# Spatial and temporal patterns of nitric oxide diffusion and degradation drive emergent cerebrovascular dynamics

**DOI:** 10.1101/836148

**Authors:** William Haselden, Ravi Teja Kedarasetti, Patrick J. Drew

**Author notes:** Correspondence (PJD).

## Abstract

Nitric oxide (NO) is a gaseous signaling molecule that plays an important role in neurovascular coupling. NO produced by neurons diffuses into the smooth muscle surrounding cerebral arterioles, driving vasodilation. However, the rate of NO degradation in hemoglobin is orders of magnitude higher than in brain tissue, though how this might impact NO signaling dynamics is not completely understood. We used simulations to investigate how the spatial and temporal patterns of NO generation and degradation impacted dilation of a penetrating arteriole in cortex. We found that the spatial location of NO production and the size of the vessel both played an important role in determining its responsiveness to NO. The much higher rate of NO degradation and scavenging of NO in the blood relative to the tissue drove emergent vascular dynamics. Large vasodilation events could be followed by post-stimulus constrictions driven by the increased degradation of NO by the blood, and vasomotion-like 0.1-0.3 Hz oscillations could also be generated. We found that these dynamics could be enhanced by elevation of free hemoglobin in the plasma, which occurs in diseases such as malaria and sickle cell anemia, or following blood transfusions. Finally, we show that changes in blood flow during hypoxia or hyperoxia could be explained by altered NO degradation in the parenchyma. Our simulations suggest that many common vascular dynamics may be emergent phenomenon generated by NO degradation by the blood or parenchyma.

## Introduction

Increases in neural activity in the brain typically are followed by the dilation of nearby arterioles^1–6^ and potentially capillaries^7, 8^ (but see^9^). This relationship between neural activity and hemodynamic signals is known as neurovascular coupling (NVC). The dilation of these vessels lowers the local vascular resistance, leading to a local increase in blood flow and oxygenation that is the basis of many brain imaging techniques^6, 10^. The maintenance of adequate coupling is thought to play a critical role in brain health^11^. In some cases, the vasodilation driven by increased neural activity is followed by a post-stimulus decrease in blood volume and flow below the pre-stimulus baseline, known as the post-stimulus undershoot^12–14^. This post-stimulus undershoot is not always observed, and its origin is not understood^15^. In addition to the post stimulus undershoot, arteries show spontaneous oscillations in diameter in the 0.1-0.3 Hz range, known as vasomotion^16–22^, whose origin is not understood. Thus, in addition to dilations linked to increases in neural activity, cerebral arterioles show a wide range of dynamic behaviors.

Multiple signaling pathways have been implicated in coupling neural activity to increases in blood flow^23^. Signals from astrocytes^8, 24–26^ and neurons^27–32^ are both thought to contribute to driving neurovascular coupling. One pathway implicated in neurovascular coupling is nitric oxide (NO)^33–35^. NO is vasoactive^36^ and affects neural excitability as well^37^. NO diffuses through aqueous and lipid mediums^38, 39^, allowing for temporally and spatially complex signaling dynamics^40–42^. NO is produced by three types of nitric oxide synthases (NOS)^43, 44^. The neuronal NOS (nNOS or NOS1) subtype of NOS is expressed by neurons^45^, while endothelial cells express endothelial NOS^46^ (eNOS or NOS3), and synthesis of NO by both enzymes is coupled to intracellular calcium^47^. An inducible, non-calcium dependent form of NOS is found in macrophages and other cells^48^ (iNOS or NOS2), and is not found in the healthy brain. NO activates guanylyl cyclase (GC) in nearby cells to produce a rise in cGMP^49^ and elicit vasodilation^38, 50–53^. Despite the importance of NO in neurovascular coupling, *in vivo* measurements of NO levels in the brain have remained elusive^54^. Direct physiologic measurements of NO concentrations in tissue is confounded by recruitment of iNOS during injury and the non-specificity of probes^55^ which may account for the large range in NO concentration reported in the literature^54^. At high concentrations, NO will block respiration in mitochondrial cytochrome c oxidase (CcO), and result in cellular damage from inhibited respiration and free radical formation^56, 57^. Because of this toxic effect on mitochondrial respiration, there will be an upper bound on NO levels in the healthy brain. Understanding the role of NO in neurovascular coupling is a topic of ongoing research^8, 11, 34, 54, 58–67^. Modulation of NO production within the physiological range has been shown to precede vascular responses^60^ and modulation of NO availability alters baseline vessel diameter^35, 62^. Inhibition of NO production blunts or abolishes the hemodynamic response^33, 34^ and causes reduction in baseline blood flow^35^. NO has been speculated to play a modulatory rather than a direct role in neurovascular coupling because adding a NO donor while inhibiting NOS can rescue the hemodynamic response^62^, though NO has a role in increasing neuronal excitability^68–72^, making the interpretation of these results difficult.

NO levels will depend not only on the dynamics of NO production, but also the degradation rate. In the tissue, NO degradation is proportional to the partial pressure of oxygen so levels of NO will tend to inversely vary with tissue oxygenation^73, 74^. However, the majority of NO is scavenged by hemoglobin in the blood and can do so a thousand-fold faster than the surrounding tissue^73, 75–78^. Because NO reacts with hemoglobin at much higher rates than the tissue, the hemoglobin present inside a vessel plays an appreciable role in shaping NO concentrations at the smooth muscle where it acts. Under normal conditions, most hemoglobin in the blood in confined to red blood cells, with low levels in the plasma. Due to fluid dynamics^79–81^, red blood cells will be excluded from the few micrometer-thick cell free layer next to the endothelial cells, providing a measure of spatial separation between the region of high NO degradation and the smooth muscles. However, if hemoglobin levels in the plasma rise (due to pathology or other processes)^82–89^, this will greatly increase the degradation rate of NO in the plasma, leading to decreased NO levels in the smooth muscle^83, 90–92^. NO’s diffusive properties and known reaction rates lend themselves to computational approaches to understanding NO signaling^38, 59, 75, 78, 93–98^. While there have been detailed and informative models of NO signaling from endothelial cells^59, 91, 96, 99, 100^ showing that the size of the arteriole^75^ and properties of the blood^96^ are vital components to understanding NO signaling, the insight from these models that the spatial location of blood plays an important role in the degradation of NO has not been applied to neurovascular coupling or in a dynamic setting.

Intriguingly, *in vitro* experiments have shown that NO released by endothelial cells can depolarize axons^67^, and flow changes in vessels can alter interneuron activity^65^, potentially providing a mechanism by which vascular cells can modulate neural activity. The idea there is bidirectional signaling between neurons and the vasculature (‘hemo-neural’ hypothesis^101, 102^) has been put forward, though there is no definitive candidate mechanism. Signaling through NO-dependent pathways is a possible mechanism for the hemo-neural coupling, as NO is known to affect neural excitability and vasodilation, an the amount of blood present could impact NO levels in the parenchyma through scavenging.

In order to understand how neuronal sources of NO communicate with the vasculature, we simulated NO production around a penetrating arteriole. In this model, the diameter of the vessel was dynamically dilated in response to the levels of NO present in the smooth muscle. We found that the sources of NO needed to be close to the arteriole to prevent inhibition of mitochondrial respiration. The increased amount of hemoglobin present during dilation greatly increased the removal of NO, which drove arteriole dynamics such as vasomotion and a post-stimulus undershoot. The concentration of plasma free hemoglobin in the blood was able to alter these vasodynamics. NO was able to function as an oxygen sensor in our model because its rate of removal in the parenchyma is dependent on the partial pressure of oxygen in the tissue. Finally, simulations imposing increases in vessel diameter when NO production rates were not varied resulted in a decrease in NO levels in the parenchyma, suggesting a potential mechanism for hemo-neural coupling. These results suggest that the diffusion and degradation of NO can drive emergent vascular dynamics.

## Results

In the cortex, penetrating arteries enter into the parenchyma perpendicular to the pial surface, and supply blood to a cylindrical volume of brain tissue approximately a hundred microns in radius^103–107^ (**Figure 1A**). The arteriole is surrounded by a layer of smooth muscle, and most of the branches off the arteriole are found in the deeper layers of cortex. The diameter of the penetrating arteriole is an important (but not the only) regulator of local blood flow^8, 108, 109^. Because the geometry of the vasculature is complex and variable^110, 111^, we simplified the geometry to a single penetrating arteriole surrounded by a cylinder of neural tissue (**Figure 1B**). The model consisted of five layers, the red blood cell (RBC) core, the cell free layer, the endothelial cells, smooth muscle and the parenchyma (**Figure 1C,D**). Each layer has an associated NO production and degradation rate, and NO is produced in both the endothelial cell layer and the parenchyma. The thicknesses of the RBC core and cell-free layer were specified for each diameter according to fits taken from empirical data^79, 80^. Unless otherwise specified, the concentrations of NO and oxygen were calculated using Fick’s equation and the Krogh model respectively. The parenchyma was treated as a nearly incompressible linear elastic solid, but the total NO production rate was held constant during dilation-induced deformations. We determined if the flow of blood will have an impact on smooth muscle NO levels using a model that includes the transportation of NO by movement of blood (see Methods). While the flow of blood will cause a convection of NO downstream, at physically plausible blood flow velocities the impact on NO concentrations is miniscule (**Figure 2–figure supplement 1**), consistent with a high Damkohler number (ratio of diffusion to convection)^75, 99, 112^. The lack of an appreciable effect of convection by the blood on NO levels allowed us to ignore convection, greatly simplifying our simulations.

**Figure 1.**
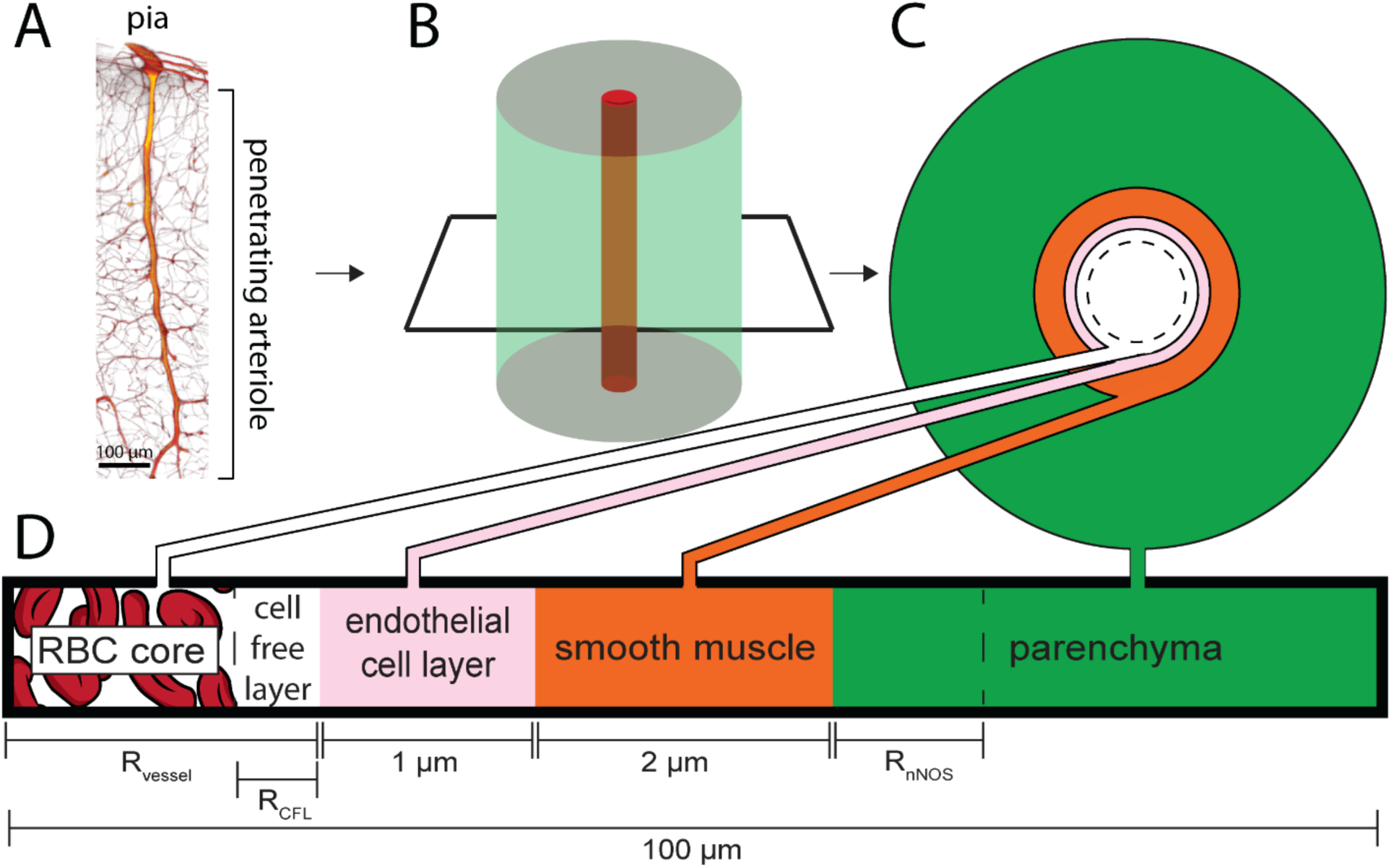
Schematic of the model. A) 3D reconstruction from serial 2-photon tomography of a penetrating arteriole. Penetrating arterioles are oriented perpendicularly to the pial surface. B) Simplified geometry used in the simulation where the penetrating arteriole is modeled as single arteriole surrounded by a cylinder of parenchymal tissue. C & D) Locations and thicknesses of the tissue domains in the model. At the center are red blood cells (RBC core). The cell free layer is a thin layer of plasma lacking red blood cells immediately adjacent the endothelial cell layer. Both the RBC core and the cell-free layer size are dynamically changed when the vessel dilates or constricts. The endothelial cells and smooth muscle make up the arterial wall, and the vessel radius is taken to be the distance from the center of the vessel to the inner wall of the endothelial cells. Outside the smooth muscle is the parenchyma, composed of neurons, glia and extracellular space. The simulated tissue cylinder is 100 µm in diameter. The thickness of the NO-synthesizing portion of the tissue (R_nNOS_), vessel diameter (R_vessel_) and the size of the cell free layer (R_CFL_) were parametrically varied.

### Effects of vessel size and NO production location on smooth muscle NO concentration

We first asked how the spatial arrangement of NO production relative to the arteriole and the size of the arteriole impacted the concentration of NO in the smooth muscle. We explored three different spatial profiles of NO production (**Figure 2A**). Early models of NO diffusion dynamics assumed homogenous NO production in the tissue^38, 113^, which we refer to as the ‘uniform’ condition. However, there is anatomical evidence that nNOS-expressing neurons and their processes may be concentrated around arterioles^32, 114, 115^ (**Figure 2A**, proximal). In the proximal geometry, all NO was produced within 2 µm of the smooth muscle^114^. We also considered an intermediate case, which we refer to as the ‘regional’ geometry. In this case, NO production is higher within 50 µm of the vessel^32^. In the uniform case, NO is produced uniformly throughout the parenchyma. We emphasize that we do not mean for these geometries to be detailed reconstructions of the actual NO production, but rather exemplars that allow us to understand the role of the spatial distribution of NO production in neurovascular coupling. We parametrically varied NO signaling for each combination of arteriole diameter, NO production and NO production location (**Figure 2D**) and evaluated their ability to signal the arteriole by the effective activation of guanylyl cyclase (GC) in the smooth muscle (**Figure 2C**). To match a given concentration of NO in the smooth muscle for a given geometry, the rate of NO production was varied. This is shown in **Figure 2B**, where the NO production rate required to reach 50% of the maximal activation of GC in a 40 µm diameter arteriole (outlined with a box in **Figure 2D**) was 0.02 M/s for the proximal geometry, 0.05 M/s for the regional geometry and 0.056 M/s for the uniform geometry.

**Figure 2.**
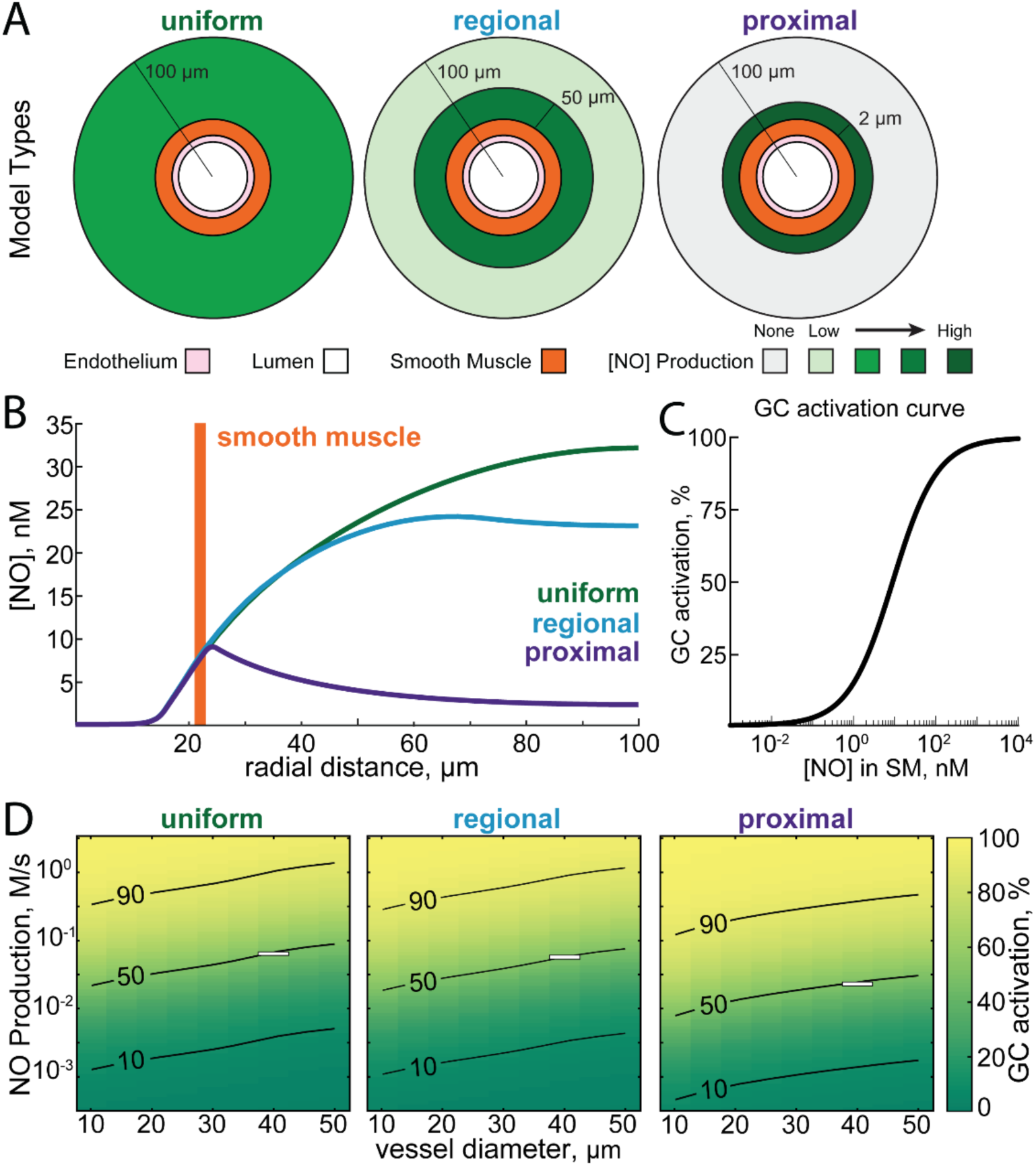
Impact of the location of NO production on NO concentration in the smooth muscle and tissue. A) Schematic showing the three simulated distributions of neuronal NO production relative to the vasculature. In the uniform model, neuronal NO-production is uniformly distributed through the parenchyma. In the regional model, there is a higher density of neuronal NO production near the vessel (within 50 μm)^32^. In the proximal model, all neuronal NO is produced within 2 micrometers of the arterial wall^32, 114, 115^. B) Plot of NO concentrations versus radial distance for each of the three models where the production rates have been chosen to yield equal concentration of NO in the smooth muscle layer (NO production rate for proximal: 0.02 M/s; regional: 0.05 M/s; uniform: 0.056 M/s). Note that the concentration of NO in the parenchyma is very different for each of these models. C) Relationship between [NO] in the smooth muscle and percent of maximal guanylyl cyclase activity in the model, based on experimental data in^50, 184, 209^. D) Plot showing percent of maximal guanylyl cyclase activation in the smooth muscle as a function of the NO production rate and vessel diameter in each of the three geometries. Superimposed curves show 10, 50, and 90% of maximal guanylyl cyclase activation. White boxes show the NO production rates and vessel diameters shown in B.

We find that when holding the rate of NO production constant, the size of the vessel had an impact on the concentration in the smooth muscle. This can be seen by the upwardly sloping contour lines in all of the NO production geometries (**Figure 2D**). If there was no size dependence, these contours would be horizontal. This size dependence was due to the higher degradation rate of NO in the hemoglobin rich portion of the blood relative to the degradation rate in the tissue. As arteriole diameter increases, more hemoglobin is present and more NO will be removed such that a higher production rate of NO is required to maintain the same concentration of NO in the smooth muscle. This parallels the experimental observation that the dilation of a vessel, as measured as a percentage of its baseline diameter, is inversely related to its resting size^1, 116, 117^, suggesting that degradation of NO by hemoglobin may contribute to size-dependence of arteriole reactivity. We explore the impact of vessel-size dependence of NO degradation in our dynamical models of dilation below.

We also find for a given concentration of NO in the smooth muscle, the different NO production geometries show markedly different concentrations of NO in the parenchyma (**Figure 2B**). This is because NO is not only degraded in the blood but also in the tissue (albeit at a substantially lower rate). The further the NO must diffuse to reach the smooth muscle, the larger the fraction that will be degraded before reaching its target. This means that the concentration of NO at a distant source (the parenchyma in the uniform model) must be higher than for a closer source (the proximal model). This high concentration of NO in the brain tissue for the uniform and regional production models can have adverse effects on mitochondrial respiration when oxygen levels fall, which we explore below.

### Impact of NO levels on mitochondrial respiration

We set out to determine the impact the spatial pattern of NO production has on mitochondrial respiration. High levels of NO are toxic, because NO competes with oxygen for the rate-limiting enzyme in aerobic respiration, cytochrome c oxidase^56, 118^. At very high levels of NO and low levels of oxygen, the reaction of NO with cytochrome c oxidase can be irreversible^119^. This inhibition of mitochondrial respiration by NO puts an upper limit on the levels of NO present in the healthy brain. Using the NO concentration profiles calculated above, combined with peri-arterial oxygen profiles derived from *in vivo* oxygen measurements using phosphorescent oxygen probes^120–123^, we calculated the cytochrome c oxidase activity as a function of distance from the simulated penetrating arteriole (**Figure 3A**). Close to the artery, the capillary density is low, and oxygenation of tissue is largely supplied by the artery^124^. As respiration depends on oxygen tension, the respiration rate will fall with distance from the vessel. However, this only becomes an issue for regional and uniform models of NO production. At levels of NO production that drive high levels of guanylyl cyclase activation in the smooth muscle, the combination of high NO levels and low levels of oxygen will result in irreversible inhibition of mitochondrial respiration in the tissue distant from the vessel. A parameter sweep of NO production rates (expressed as guanylyl cyclase activation in the smooth muscle) and vessel size shows that for both the regional and uniform models, high levels of NO production can be toxic for an appreciable fraction of the tissue (**Figure 3B**). Though this hypoxia will be mitigated by capillaries supplying oxygen to tissue distant from the arteriole, these simulations suggests that keeping the site of NO production close to the smooth muscle may prevent tissue damage associated with high NO levels.

**Figure 3.**
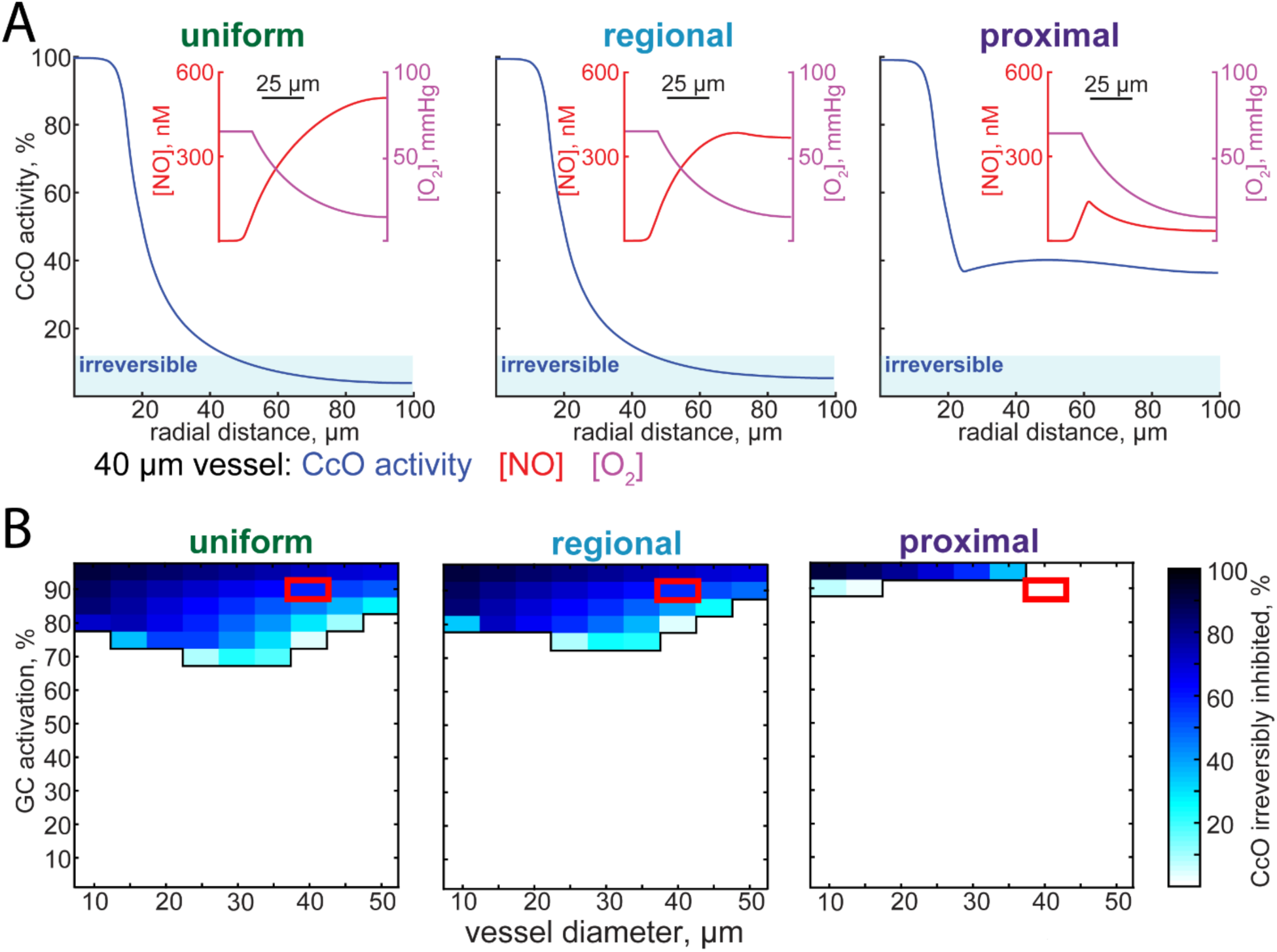
Extent of the NO inhibition of mitochondrial respiration depends on the location of NO production. A) Plots of cytochrome c oxidase (CcO) activity as a function of radial distance for the uniform, regional, and proximal models. The vessel diameter was set to 40 micrometers, and NO production rates have been set so that there is 90% of maximal GC activation in the smooth muscle. When CcO activity drops below 12.5%, CcO inhibition is irreversible^56^, and this is shown as a shaded region labelled ‘irreversible’. Insets show oxygen and NO concentrations as a function of radial distance. Oxygen concentration curves match *in vivo* measurements^120, 121^. B) The fraction of tissue irreversibly inhibited as function of various NO production levels and vessel diameters for each of the three different NO production geometries. Red boxes indicate simulations plotted in (A). Note that for the regional and uniform NO production geometries, CcO inhibition becomes an issue at a wider range of NO production levels. For the proximal production case, inhibition of respiration by NO only occurs at the highest levels of NO production.

### Biphasic hemodynamic responses from increased NO removal by blood during vasodilation

A larger arteriole degrades more NO than a smaller one, enough to alter NO levels appreciably in smooth muscle at steady state. We then investigated whether a similar process could occur during vasodilation and what impact it would have on vasodynamics. We moved to a dynamic model, in which the concentration of NO in smooth muscle dynamically dilated the vessel (**Figure 4–figure supplement 1**). An important parameter in these simulations is the sensitivity of the dilation to changes in GC activation, captured in our simulations in the parameter ‘m’, (which will have the units of 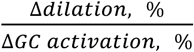, see Methods). The sensitivity of arteries to NO is known to vary^50, 125–128^, and the larger m is, the more sensitive the artery is to changes in NO concentrations. Empirically, studies suggest m is in the range of 1-5, with m=5 giving dilations comparable to the largest stimulus evoked dilations in awake animals^1, 117, 129, 130^. The key interaction in this model was that the dilation of the arteriole caused an increase in the local hemoglobin concentration via an increase in the size of the red blood cell-containing ‘core’ (RBC core) (**Figure 4–figure supplement 1, Figure 4B**). This increase in hemoglobin would in turn cause an increase in NO degradation, which functions as a delayed negative feedback on NO levels in the smooth muscle. The dilation will be delayed relative to the increase in NO production due to diffusion time, and the latency of the signal transduction cascade transducing the elevation of NO levels in the smooth muscle into relaxation. We wanted to understand if this separation of timescales could produce vasodynamics like those seen *in vivo*. For all subsequent simulations we use the proximal NO production geometry.

**Figure 4.**
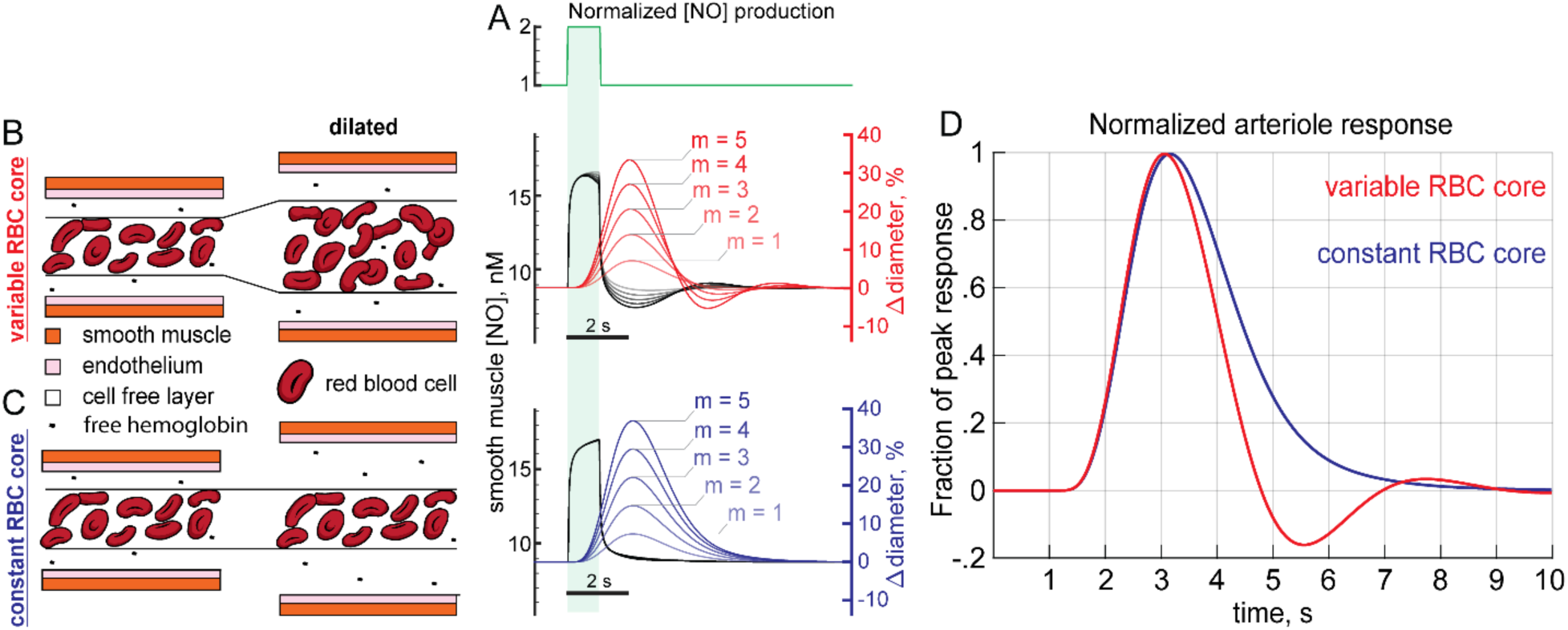
Dynamical model of NO-induced dilation shows a post-stimulus undershoot. A) NO production in the proximal model was increased 100% for 1 second in these simulations with a baseline NO production set at EC_50_ in the smooth muscle. B) Arteriole dilation increases the supply of RBCs in the ‘physiologically realistic’ variable RBC model. The arteriole dilates (red) in response to increased NO in the smooth muscle (black); however, after NO production returns to baseline the arteriole is still dilated. The dilated arteriole can accommodate more RBCs which depletes NO below baseline. The depletion of NO concentrations below baseline is reflected in a corresponding post-stimulus constriction. Five different sensitivities to GC 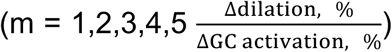 are shown. C) Dilation in the constant RBC core case does not increase RBC supply or the degradation rate of NO. When vasodilation does not increase NO consumption, NO concentrations do not fall below baseline and no post-stimulus constriction occurs. D) Dilations for 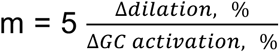 rescaled to the same height showing the relative size of the post-stimulus constriction in the variable RBC core case while none is present if the RBC core is held constant during vasodilation.

We first simulated the effects of a transient increase in NO production, similar to what would be generated in response to a brief elevation of neural activity in response to a stimulus. The effects of a stimulus were implemented by doubling parenchymal NO production for 1 second (**Figure 4A**). When the increased arteriole diameter elevated NO scavenging by increasing the amount of hemoglobin (**Figure 4B**), NO concentrations in the smooth muscle dropped below baseline during vasodilation (**Figure 4B**, black), even though there is no corresponding decrease of NO production below baseline (**Figure 4A**). The drop in NO concentration in the smooth muscle results in a post-stimulus undershoot (**Figure 4B**, red), reminiscent of the canonical HRF seen *in vivo*^1, 28, 131^. We hypothesized that this undershoot was driven by the increased hemoglobin in the vessel that would naturally take place when the vessel dilated. To test this hypothesis, we performed simulations where the RBC core was kept at a constant diameter when the arteriole dilated (**Figure 4C**). Without the increase in NO degradation mediated by an increase in hemoglobin, the post-stimulus undershoot was not observed (**Figure 4C**, blue). To better visualize the differences between the two conditions, we plotted the two responses together (**Figure 4D**). The (physically realistic) variable core model shows a clear undershoot, (**Figure 4D**, red) while the constant core model does not (**Figure 4D**, blue). The variable core model could generate arterial dilation dynamics qualitatively similar to those seen in awake mice in response to sensory stimulation^1^ (**Figure 4–figure supplement 2**). Together, these suggest that the increased NO scavenging in the RBC core during vasodilation can be a contributing factor to the post-stimulus undershoot in arterial diameter.

### Interplay of vasodilation and NO degradation can generate vasomotion oscillations

We then sought to quantify the effects of an increase in NO scavenging accompanying dilation on arteriole dynamics. The relationship between a stimulus or neural activity and the vascular dynamics is captured by the hemodynamic response function^131^ (HRF). The HRF is a linear kernel that low-pass filters neural activity into a change in vessel diameter. This kernel can be easily extracted from the response (in this case, artery diameter) to a spectrally white input^117, 132, 133^ (in this case, NO production linked to neural activity). To better understand how NO degradation dynamics impact neurovascular coupling, we simulated the response of both the variable RBC core and constant RBC core models (**Figure 5G**, red & blue) to randomly varying (‘white noise’) NO production (**Figure 5G**, black). We then deconvolved out the HRF (using the modified Toeplitz matrix method^131^) (**Figure 5B,E**) from the vascular response. We found that there was an undershoot in the variable RBC core model (**Figure 5B**), but no undershoot following the dilation of the constant RBC core model (**Figure 5E**). The undershoot was driven by the decreased NO levels in the smooth muscle accompanying dilation due to the larger amount of hemoglobin in the dilated artery (**Figure 4B**), and the magnitude of the undershoot increased with increasing sensitivity to NO (**Figure 5B**). Even though the undershoot was an emergent property in the simulations, it was still linear, as the variance explained by the HRF was very high (R^2^∼0.95) (**Figure 5–figure supplement 1**). By looking at the power spectrum of the arteriole diameter changes elicited by white noise NO production we can see the frequency response properties of the system. Interestingly, the power is highest in the 0.1 - 0.3 Hz frequency band of the power spectrum of the artery diameter in the variable RBC core model (**Figure 5C**), showing that this system effectively acts as a band pass filter. This peak is reminiscent of vasomotion, a rhythmic 0.1 - 0.3 Hz oscillation in cerebral artery diameter observed in awake and anesthetized animals, *in vitro* and in humans^16–18, 131, 134, 135^. When the vasodilation does not increase NO scavenging, as is the case when the RBC core is held constant, no undershoot (**Figure 5E**) or elevation of power in the 0.1 – 0.3 Hz band were observed (**Figure 5F**). This comparison of the dynamic and constant RBC core models highlights the importance of NO degradation on vascular dynamics (**Figure 5H,I**). The effects of NO scavenging by increased hemoglobin likely work in concert with other drivers of vasomotion^136–139^ to generate these oscillation *in vivo*.

**Figure 5.**
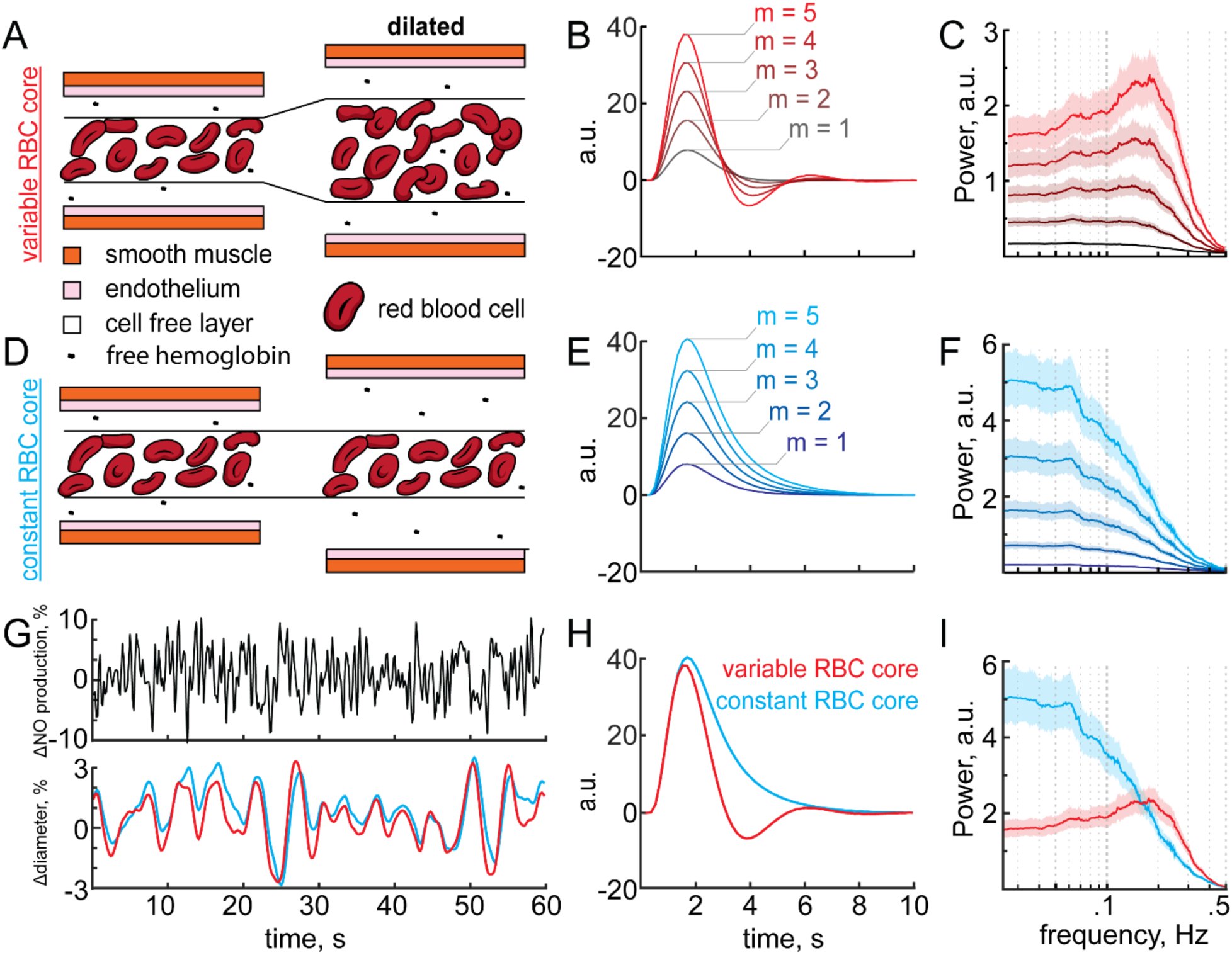
Arteriole sensitivity to NO increases the amplitude of the undershoot and vasomotion. A) Schematic of the variable RBC core model. Vasodilation increases the diameter of the RBC core and thus the degradation rate of NO in the variable RBC core model. B & C) Hemodynamic response function (B) and power spectrum (C) of the variable RBC core model from a white noise NO production rate. Note that with increasing NO sensitivity 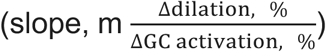 the magnitude of the undershoot and power near 0.2 Hz increases. D) Vasodilation does not increase the diameter of the RBC core and thus the degradation rate of NO does not change in the constant RBC model. E & F) No post-stimulus undershoot is present (E) and the maximum power of the constant RBC core model is at low frequencies (<0.1 Hz) (F). G) 60 second example taken from a 25 minute trial displaying NO production and resultant diameter changes from which arteriole dynamics were evaluated. H & I) Juxtaposition of variable (red) and constant (blue) RBC core models for m = 5 showing the post-stimulus constriction (H) and peak power between 0.1 – 0.3 Hz (I) in the variable RBC core case (red) while no undershoot or peak power between 0.1 – 0.3 Hz is present if the RBC is held constant during vasodilation (blue).

### Influence of plasma free hemoglobin and hematocrit on vasodynamics

Because NO is mainly degraded by the blood, we expected that changing hematologic properties such as free hemoglobin (Hgb) or hematocrit (Hct) would alter NO-mediated signaling. Hematocrit varies with sex^140^, and can be elevated by drugs^84, 141^ or prolonged exposure to high altitude^142, 143^. While NO is typically degraded by hemoglobin (Hgb) in RBCs, free Hgb in the plasma can scavenge NO up to 1,000-fold faster than in the RBC membrane^144, 145^. Under normal conditions free Hgb levels in the plasma are low, and the impact of this free Hgb on NO levels is minimal. However, plasma free Hgb can be elevated in sickle cell disease^83^, malaria^146^ or following blood transfusions^147^. Elevation of free plasma Hgb can cause cardiovascular issues^85, 148–151^ due to the increased scavenging of NO^152, 153^.

We first explored the effects of altering plasma free Hgb. Increasing free Hgb (**Figure 6A**) reduced arteriole diameter (**Figure 6B**) though depletion of perivascular NO (**Figure 6C**), consistent with *in vivo* experiments^154^. The increase in free Hgb resulted in a larger post-stimulus undershoot (**Figure 6D**), and an increase in the band-pass like properties of the arteriole (**Figure 6E**). These simulations suggest that in addition to other symptoms, elevated plasma free hemoglobin may also cause constriction of cerebral arterioles and alter the dynamics of hemodynamic responses. Increasing hematocrit resulted in decreases in baseline arteriole diameter (**Figure 6–figure supplement 1B**) and perivascular NO (**Figure 6–figure supplement 1C**) in the model. However, neither the HRF, nor frequency response properties of the vessel were appreciably affected by varying the hematocrit (**Figure 6–figure supplement 1D,E**). The lack of an effect can be attributed to the fact that even under different hematocrit concentrations the location of NO scavenging remains unchanged. However, when increasing free Hgb in the plasma, the compartmentalizing effects of the CFL is compromised and the location of NO scavenging shifts from the center of the lumen to much closer to its source^75, 97, 155, 156^. With NO being scavenged much closer to the smooth muscle, any changes to the rate of scavenging (such as increased hemoglobin during dilation) are amplified. While both hematocrit and free Hgb in the plasma contribute to determining baseline arteriole tone these simulations suggest that plasma free hemoglobin can also have a substantial effect on vasodynamics through a NO-mediated mechanism.

**Figure 6.**
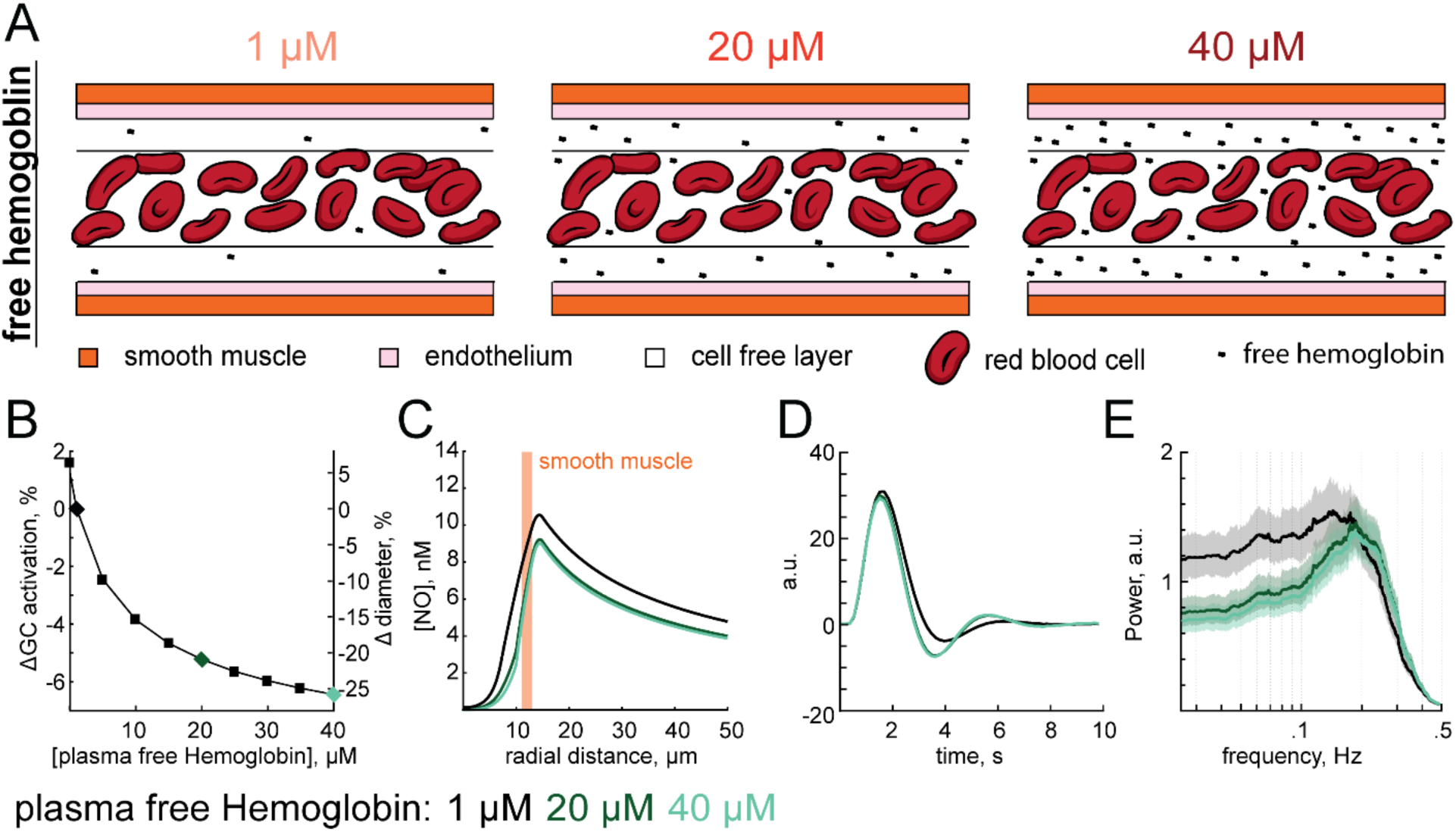
Impact of plasma free hemoglobin on vasodynamics. A) When hemoglobin (Hgb) is released into the plasma, the location of NO degradation shifts from the RBC core to the Hgb rich cell free layer. B) Increasing free Hgb constricts arterioles. The change in diameter was calculated at steady state after accounting for the change in NO degradation. Black, dark green and light green diamonds correspond to 1, 20 and 40 μM of plasma hemoglobin respectively with colors interpretations conserved from B to E. C) The shift in location of NO consumption to the cell free layer and increased reactivity of free Hgb over RBC Hgb decreases perivascular NO with diminishing returns past 20 μM. D) Increasing free hemoglobin slightly increases the undershoot in the hemodynamic response function. E) Increasing free hemoglobin reduces the low frequency power (<0.1 Hz) and strengthens the band pass properties within the 0.1 - 0.3 Hz range.

### NO can act as sensor of cerebral oxygenation

Despite the lack of an known oxygen sensor in the brain, hypoxia will dilate and hyperoxia will constrict cerebral arterioles^157–174^. These cerebrovascular responses to blood oxygenation are modulated by NO availability^157, 169, 175–179^, occur under isocapnic conditions^177, 179^ and constant pH^177^. We wanted to investigate if changes in NO consumption due to oxidative reactions in the parenchyma could contribute to hypoxia-induced vasodilation. The first order dependence of NO removal on tissue oxygen concentration^73^ would mean that NO would be degraded faster under a hyperoxic condition. Elevated oxygen concentrations could constrict arterioles by depleting perivascular NO, and conversely low oxygen could dilate arterioles by consuming less NO, effectively allowing NO to functioning as a local oxygen sensor. We tested this idea by dynamically varying the oxygen levels in the artery (**Figure 7A,B**) and looking at the resulting changes in vessel diameter (**Figure 7D**) due to changes in parenchymal NO degradation (**Figure 7C**). Using a baseline arteriole oxygen concentration of 65mmHg^120–123^, we varied arteriole oxygen tension in the range from 0 to 125 mmHg^180^. Hypoxia drove dilation, and hyperoxia drove vasoconstriction, though not with as large of magnitude (**Figure 7E**). The observation that in our model hypoxia drove a larger dilation than hyperoxia drove constriction is consistent with *in vivo* observations^166, 170, 181^. These results highlight NO’s potential to function as a local oxygen sensor by linking perivascular oxygen concentrations to vascular tone through an oxygen dependent rate of NO removal in the parenchyma.

**Figure 7.**
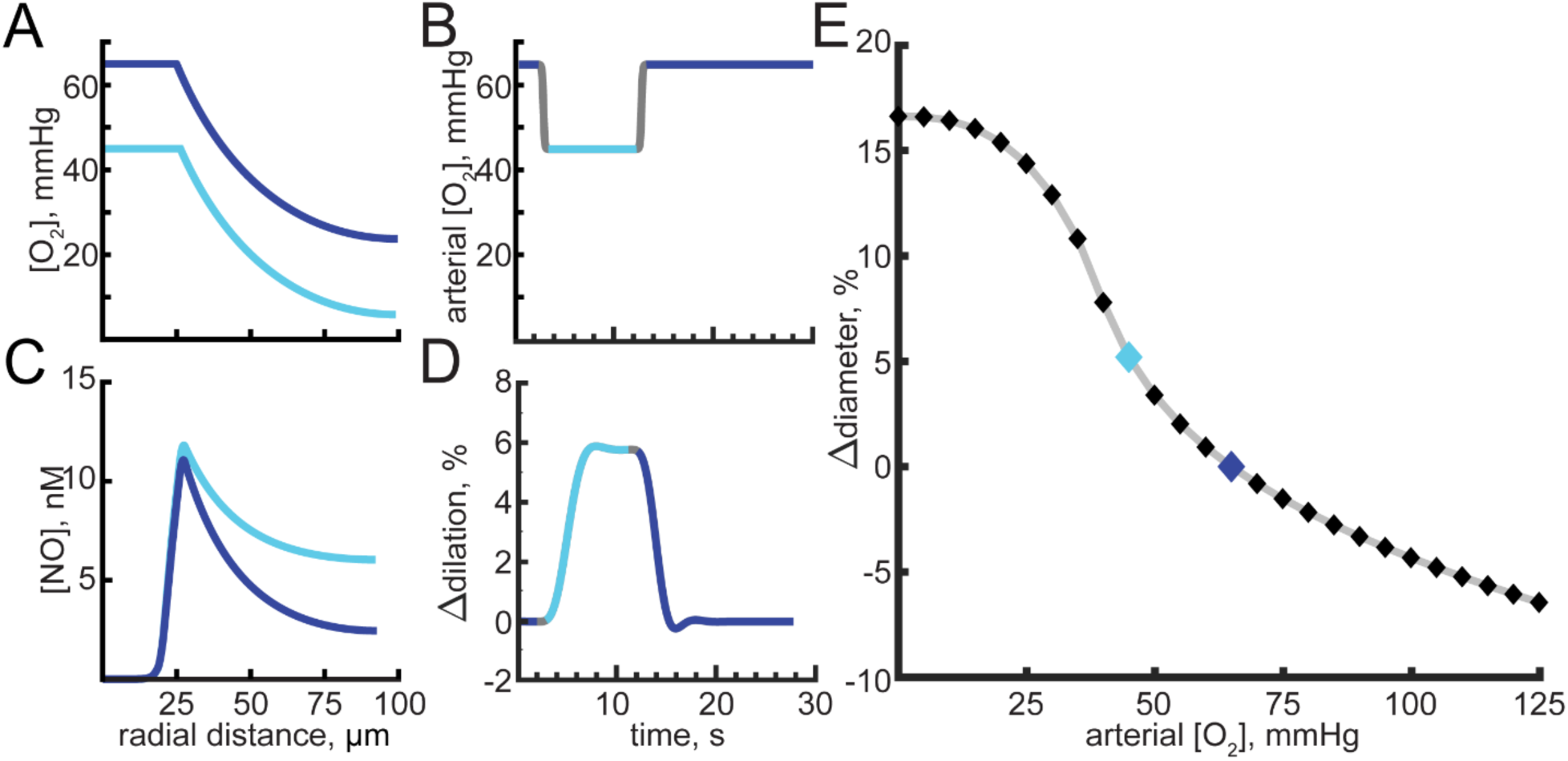
Hypoxia and hyperoxia alter NO levels and can drive vasodilation and vasoconstriction. A) Oxygen concentration as a function of distance from the arteriole center with a blood oxygen content of 65 mmHg (dark blue) or 45 mmHg (light blue). B) Time course of arterial simulated oxygen content. The oxygen concentration drops 20 mmHg for 10 seconds before returning back to 65 mmHg. Gray indicates time at which arteriole oxygen tension is changing and dark blue and light blue indicate arteriole oxygen content of 65 mmHg or 45 mmHg, respectively. C) Perivascular NO concentrations with a 65 mmHg (dark blue) and 45 mmHg (light blue) blood oxygen content. D) Arteriole response from a 10 second 20 mmHg decrease in blood oxygenation shown in (B). Arteriole sensitivity to NO is set to m = 4. E) Hypoxia increases arteriole diameter at a more rapid rate that hyperoxia. Dark and light blue diamonds correspond to blood oxygenation states shown in (A-D).

### Impact of vasodilation on parenchymal NO concentration

It has been proposed that changes in the vasculature can drive changes in neural activity^101^. As the degradation of rate of NO is greatly influenced by the amount of hemoglobin and NO levels affect neural excitability^69, 70^, we hypothesized that changes in NO concentration driven by vasodilation might be able to drive changes in NO levels of nearby neurons. In all our previous simulations, the concentration of NO in the smooth muscle has thus far changed with size of the arteriole. Therefore, we then asked how vasodilation due to other pathways^2, 8, 11, 23–26, 30, 32, 182^ will impact parenchymal NO levels. We isolated the influence of vasodilation on parenchymal NO in the model by imposing changes in arteriole diameter (**Figure 8B**) in the background of a constant NO production rate (**Figure 8A**). This vasodilation caused a decrease of NO in the smooth muscle (**Figure 8C**). Because as the vessel dilates, it slightly distorts the tissue, we looked at the parenchymal NO concentrations relative to the outer edge of the smooth muscle and adjusted for deformation, rather than from the vessel centerline. We found that vasodilation caused an appreciable drop in the NO concentrations in the parenchyma (**Figure 8D**). We then parametrically varied the sign and amplitude of the vessel diameter change and looked at the impact of these diameter changes on parenchymal NO levels. We found that dilation and constriction in a physiologically plausible range can produce substantial changes in parenchymal NO (**Figure 8E**), large enough to potentially alter neural excitability. These simulations identify a potential mechanism by which neurons can receive information about the state of nearby vessels.

**Figure 8.**
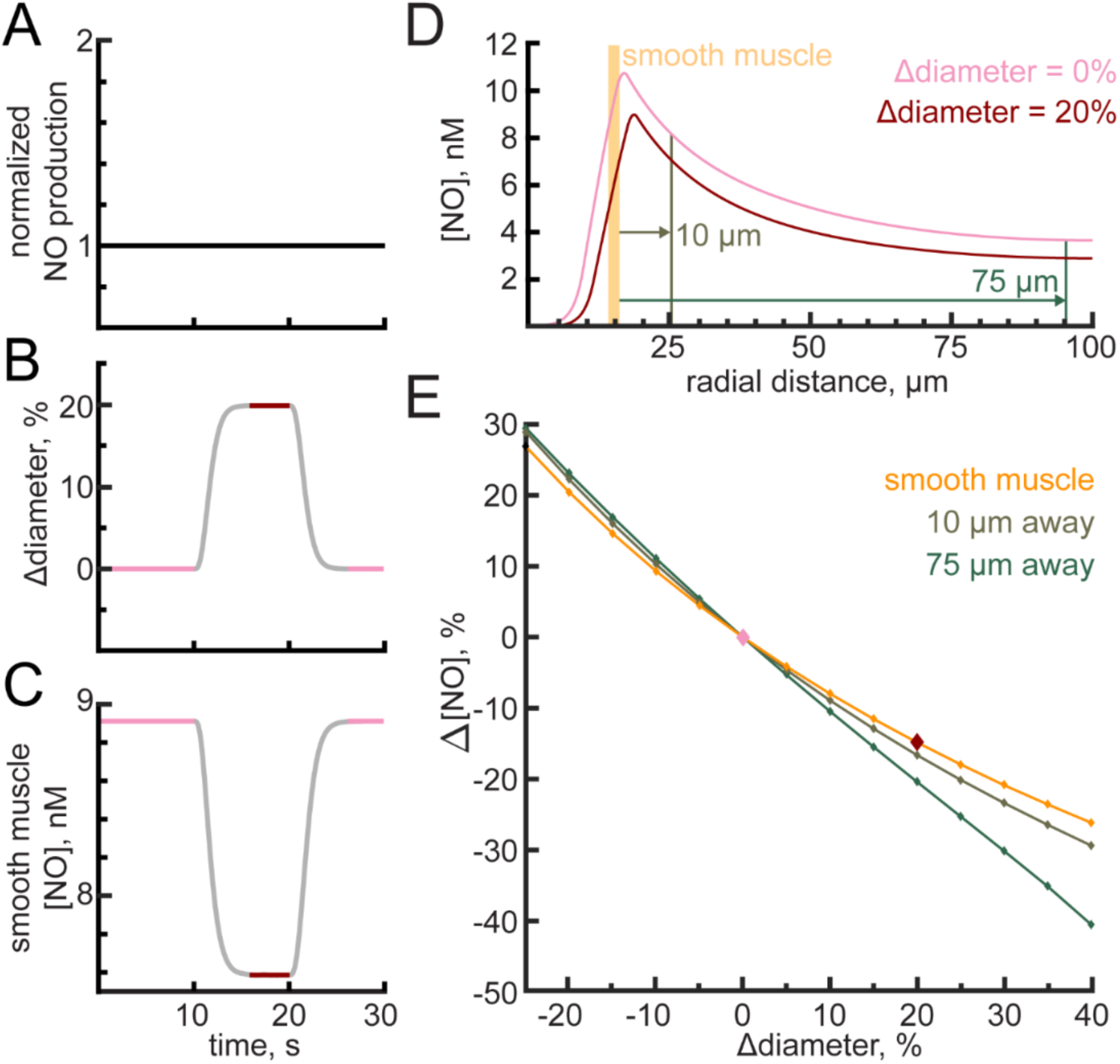
Arteriole diameter changes alter perivascular NO. A) NO production was held constant. (B) A 20% dilation of the arteriole was externally imposed. Pink indicates the pre-dilated state while red is the dilated state. Grey indicates transition times in which steady state has not yet been reached. C) NO concentrations in the smooth muscle depleted as a result of the increased arteriole diameter. D) The perivascular NO in the smooth muscle (orange), 10 μm from the arteriole wall (brown), and 75 μm from the arteriole wall (green) were all decreased following a 20% dilation. The reference position of the smooth muscle is shown in orange, and parenchymal position markers in brown and green is for the pre-dilated state. E) Perivascular NO changed during arteriole dilation and constriction even at distances up to 75 μm from the arteriole wall.

## Discussion

Computational models of diffusion have complemented experimental techniques to give us insights into how NO signals the vasculature. Pioneering work by Lancaster^38^, Wood and Garthwaite^39^, and others^59, 76, 91, 96, 97, 99, 100, 178, 183^ have demonstrated the importance of NO degradation by the blood in shaping the efficacy of NO signaling. Building upon their work, we apply the insights gained from modeling NO dynamics to neurovascular coupling. By coupling NO-dependent changes in arteriole tone and blood supply to a model of NO diffusion we are able to reproduce many of the commonly observed arteriole dynamics. These include the size dependence of arteriole dilation, vasomotion, the post-stimulus undershoot, and hypoxia-induced vasodilation. We show that in addition to the neural production of NO, consumption of NO by the blood also has the potential to modulate the hemodynamic response and that many pathologically homologous conditions may disrupt neurovascular coupling via increased NO degradation.

There are several caveats to our work. Though for simplicity we did not simulate other neurovascular coupling pathways^2, 8, 11, 23–26, 30, 32, 182^, this should not be taken to mean that NO-mediated coupling is the only (or even primary) neurovascular coupling mechanism. While there is a wide range of NO concentrations measured in the tissue^54^, the levels of NO that drive GC activation have been consistent across several studies^50, 184^. Additionally, none of the results here are sensitive to the absolute levels of NO or GC activation, as they depend on the fact that degradation of NO is much higher in the presence of hemoglobin. Finally, we used simplified vascular and neural geometries in order to gain insight into how NO production and degradation dynamics might impact neurovascular coupling. Future work that combines large-scale vascular reconstructions^110, 111^ paired with detailed mapping of neuronal cell-type locations^185, 186^ in the brain will allow the creation of more realistic NO diffusion models that may give insight into the heterogeneity of neurovascular coupling across brain-regions^187–190^.

Our simulations show that NO degradation dynamics by the blood can provide a mechanism for many experimental observations of cerebrovascular dynamics^16–18, 131, 134, 135, 154, 166, 170, 181, 191^. The combination of genetically-encoded cGMP sensors^192, 193^ combined with optogenetic stimulation nNOS-expressing neurons^194^ should allow the ideas presented here to be examined experimentally. Importantly, our simulations also suggest that hemodynamic signals in the brain do not solely depend on neural activity, but rather can be greatly modulated by normal and pathological variations in the composition of blood.

## Methods

Simulations were performed in COMSOL (COMSOL Multiphysics: partial differential equations (pde) module version 5.2, Burlington MA), with a LiveLink to Matlab (version R2018b, Mathworks, Natick MA) to provide control of dynamic variables. Simulation outputs were analyzed in Matlab. A 400 µm long penetrating arteriole was modeled in the center of a 100 µm radius cylinder of parenchymal tissue with a zero flux boundary condition. Calculations were simplified by taking advantage of radial symmetry and assuming no concentration gradients in the circumferential (θ) direction. Axial gradients of NO did not play a role unless convection was considered. All domains were assumed to have homogenous properties.

### Overview of model formulation and governing equations

In addition to diffusive movement of NO, the flow of blood could add a convective component to the movement of NO. To determine whether convection of NO driven by the flow of blood plays an appreciable role in NO dynamics, we simulated fluid flow in the vessel lumen in a full 3-D model and examined the impact of blood movement on NO concentration in the smooth muscle (**Figure 2–figure supplement 1**). For flows in the physiological range, there was negligible effect of blood flow on the NO concentration in the smooth muscle. This result is consistent with the high Damkohler number (ratio of diffusion to convection) calculated in previous models of NO diffusion^75, 99, 112^. This allowed us to simplify our model for further simulations by assuming a negligible effect of convection on NO diffusion and not simultaneously model blood velocity profiles. Note that the COMSOL files also contain the ability to include convective flow calculations by specifying a non-zero pressure difference if desired (parameter: press1 [Torr]) (see data availability).

To investigate how NO scavenging by the blood sculpts hemodynamic responses we modeled the interaction at the level of a penetrating arteriole supplying blood to a region of the parenchyma (**Figure 1**). NO production rates in the parenchyma and degradation rates in the blood (**Table 1**) were used in a diffusion model to predict the hemodynamic response using the quantity of NO reaching the smooth muscle (**Figure 2C**). We generated a finite element model with this cylindrical geometry in COMSOL. The finite element model was divided into five domains: a red blood cell-containing ‘core’ (RBC core), a cell free layer, an endothelial cell layer, a smooth muscle layer, and parenchymal tissue. Each domain had their respective rate of production or degradation of NO (**Table 1**). NO was free to diffuse according to Fick’s law.

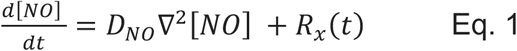

**Table 1.**
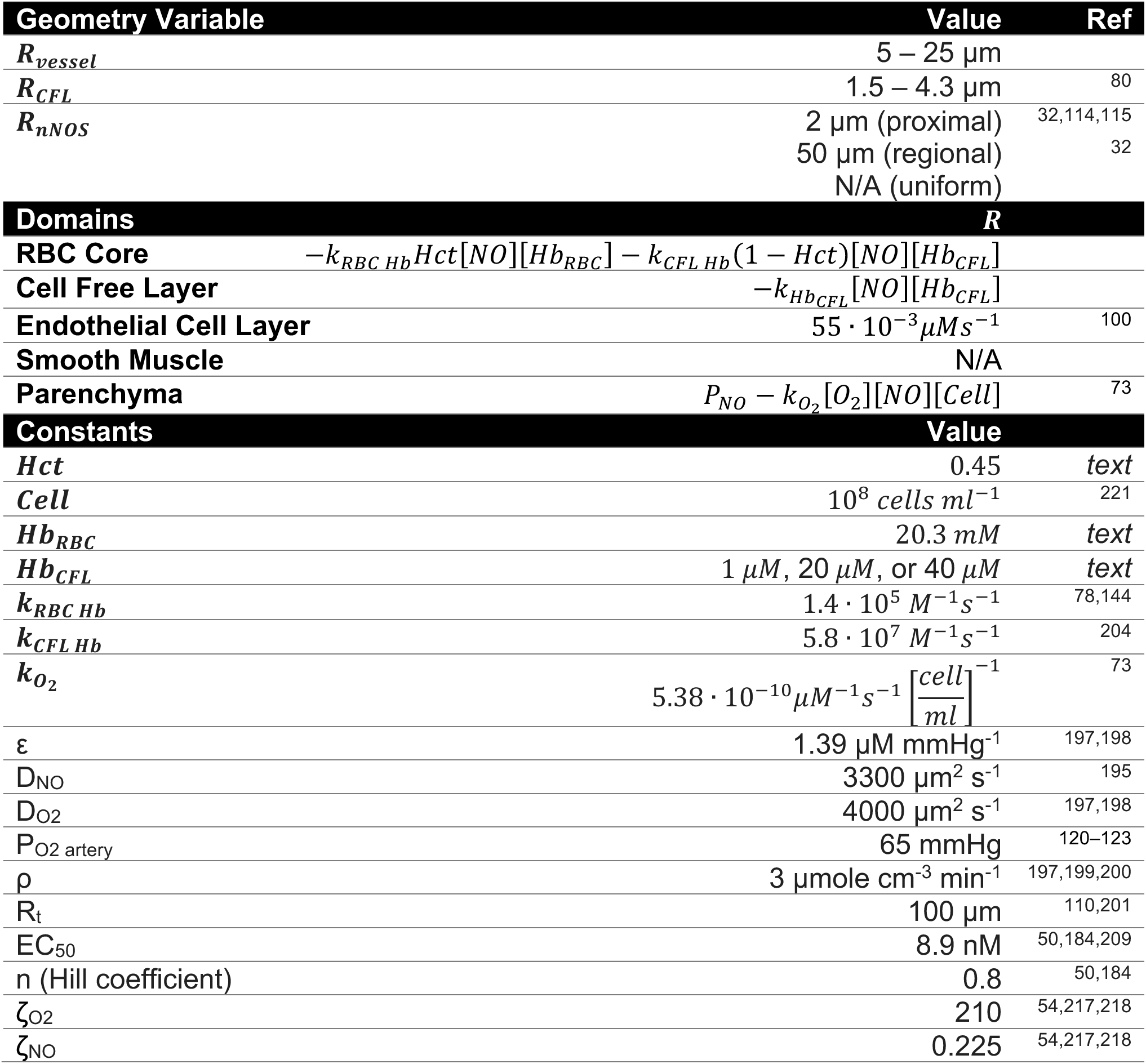
Simulation Parameters

Where [NO] is the concentration of NO at any given point in space, D_NO_ is the diffusion coefficient of NO (3300 µm^2^ s^-1^)^195^, and R_x_(t) is the time dependent degradation or production rate of NO unique to each domain (**Figure 1D, Table 1**).

Perivascular oxygenation was estimated using the Krogh model or Fick’s diffusion equation with oxygen. Luminal oxygen concentration was set to 65 mmHg^120–123^ and oxygen in the parenchymal tissue set to have a lower bound of 10 mmHg^121^. The Krogh model is a solution to radially symmetric oxygen diffusion from a cylinder (blood vessel) at steady state^196^, and is given by the equation:

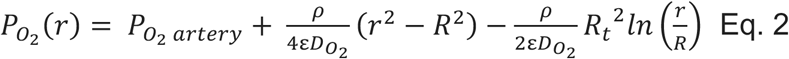

Where P_O2 artery_ is arteriole oxygen content in mmHg, D_O2_ is the diffusion coefficient of oxygen in water (4000 µm^2^ s^-1^)^197, 198^, r is the distance from the arteriole, R is the radius of the arteriole, R_t_ is the diameter of the tissue cylinder (100 µm), ε is the tissue oxygen permeability (ε = 1.39 µM mmHg^-1^), and ρ is the cellular metabolic rate of oxygen consumption (CMRO_2_) in the parenchyma, taken to be 3 µmole cm^-3^ min^-1^, as CMRO_2_ in the awake state is double that under anesthesia^197, 199, 200^. For simulations where oxygen levels change rapidly (**Figure 7**), we explicitly modelled the diffusion of oxygen from the lumen into the parenchyma with Fick’s equation:

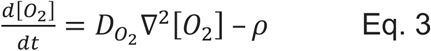

Where [O_2_] is the concentration of oxygen at any given point in space. The average distance to the nearest penetrating artery from any point in the parenchyma is of order 100 µm^110, 201^, so we modeled NO and oxygen diffusion into the parenchyma up to 100 µm from the arteriole with a repeating boundary condition (see Methods: Parenchyma). The model was initiated at steady state 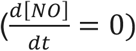 before implementing time-dependent ^c^hanges in NO production, R_x_(t). While there is disagreement as to the levels of NO in the brain^54^, the NO concentration dependence of guanylyl cyclase activity is relatively well characterized^67, 184^ and can be used to estimate the extent of vasodilation (see Methods: Smooth Muscle).

### Red Blood Cell core

Red blood cells (RBCs) are not distributed homogenously in the vessel, they cluster in the center (core) of the vessel and are excluded from the volume close to the endothelial cells^81, 202, 203^. NO entering the RBC core region is heavily scavenged by the hemoglobin contained in RBC. The rate of NO scavenging by the RBCs was obtained by multiplying the rate of NO and RBC hemoglobin interaction (k_RBC Hb_ = 1.4 • 10^5^ M^-1^s^-1^)^78, 144^ with the hemoglobin concentration in a single RBC (20.3 mM), and the core hematocrit^78^ was taken to be 0.45 unless otherwise specified. Additionally, free hemoglobin in the plasma occupying the spaces between the RBCs can also contribute to NO scavenging. Free hemoglobin is limited in the plasma (∼1 µM) compared to hemoglobin contained in RBCs, but has a much higher reaction rate with NO (k_CFL Hb_ = 5.8 • 10^7^ M^-1^s^-1^)^204^. The plasma component of NO degradation in the RBC core was calculated by multiplying the fraction of plasma (1-Hct) with the rate of NO and hemoglobin interaction in the plasma (k_CFL Hb_), and the concentration of hemoglobin in the plasma which is modulated in the model to be 1, 20, or 40 µM. The total degradation rate of NO in the RBC core was assumed to be homogeneous, and was taken to be the sum of the scavenging from RBCs and plasma components:

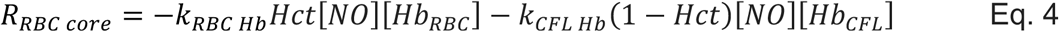

Detailed equations and parameters can be found in Table 1.

### Cell Free Layer

The cell free layer (CFL) is a layer of blood plasma between the RBC core and endothelial cells. The CFL influences NO signaling by providing a region of reduced NO degradation that increases the concentration of NO in the smooth muscle^156, 203, 205^. The thickness of the CFL can be accurately predicted given a known vessel size and blood hematocrit^80^. For 10 – 50 µm arterioles, the CFL thickness is a quantitative function of vessel diameter and hematocrit^79, 80^. The scavenging rate of NO in the cell free layer is the product of the rate of NO and hemoglobin interaction in the plasma (k_CFL Hb_ = 5.8 • 10^7^ M^-1^s^-1^)^204^:

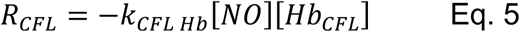

^T^he concentration of plasma free hemoglobin ([Hb_CFL_]) in the CFL was modulated in the model to be 1, 20, or 40 µM. Detailed equations and parameters can be found in Table 1.

### Endothelial cell layer

NO is not only produced from nNOS in the parenchyma but also from eNOS contained in endothelial cells. The contribution of NO from the endothelial cell layer is thought to be much smaller than parenchymal sources^78, 206^, but was still accounted for in the model by assuming a constant production rate of 55 • 10^-3^ µM s^-1^ in the 1 µm thick ring between the lumen and smooth muscle^100^.

### Smooth muscle

Upon entering the smooth muscle, NO activates guanylyl cyclase (GC) to induce vasodilation via increased cGMP production^50, 53, 207^. The relationship between NO concentration and GC activation or vessel relaxation can be described by the Hill equation with a NO half-maximal excitatory concentration (EC_50_) between 3 and 10 nM and a Hill coefficient near 1^50, 184, 208, 209^. For our model, we used an EC_50_ of 8.9 nM and a Hill coefficient of n = 0.8^50, 184, 209^ to calculate the activity of GC_f_ as a function of the average concentration of NO in the smooth muscle, [NO]:

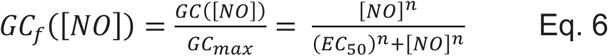

The sensitivity of an arteriole to NO can be modulated^127, 209^. In order to account for an arteriole’s ability to become sensitized or desensitized to NO, we kept changes in vessel size relatively low (±5%) when investigating vasodynamic properties and assumed a linear relationship between GC activation and vasodilation within this range. The slope of the relationship between GC activation and vasodilation was denoted by the variable 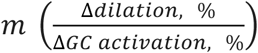 which was varied between 1 and 5 in our model.

The dilatory response following brief sensory stimuli usually peak after 1 - 2 seconds^1, 131, 133, 210^ and can be mathematically described by the convolution of the hemodynamic response function (HRF) with the stimulus. The HRF is typically modeled by fitting with a gamma distribution function^14, 211^. In some cases, in order to capture the post-stimulus undershoot the HRF is modeled as a sum of two gamma distributions, a positive one with an early peak to capture the stimulus-induced dilation, and a slower negative one to generate a post-stimulus undershoot^14, 212^. Because NO is a vasodilator and increases in GC activation are accompanied with increases in vessel diameter, we modeled the response of the vessel to NO using a single gamma function matched only to the positive component of the HRFs observed *in vivo*^131^:

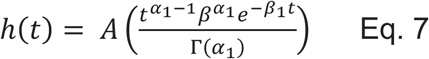

Where α_1_ = 4.5, β_1_ = 2.5, t is time and A is the amplitude which was normalized such that 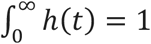^T^he predicted diameter was calculated in Matlab and transmitted to COMSOL with Matlab Livelink to dynamically adjust vessel diameter (**Figure 4–figure supplement 1**):

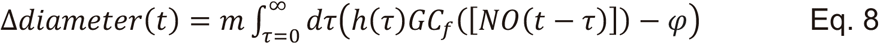

The fractional change in diameter of the arteriole was the deviation of the convolution of the HRF (**Eq. 7**) and past fractional GC activity (**Eq. 6**) from its initial state (φ = GC_f_([EC_50_])) multiplied by the sensitivity of the arteriole to NO 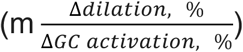. This convolution was performed at each time step so that COMSOL could recalculate Fick’s diffusion equation given the new vascular diameter. Because a larger arteriole will supply more hemoglobin which scavenges more NO this created a dynamic model in which vasodilation was linked to changing NO degradation via a changing vessel diameter.

### Parenchyma

NO is both produced and degraded in the parenchyma, although the rate of NO degradation within this region is much lower than the degradation rate of NO in the lumen. NO diffusion into the parenchyma was modeled with a reflecting (no flux) boundary condition at the radial boundaries of the simulated domain. Parenchymal NO production was geometrically varied between three models: uniform, regional, and proximal. In the uniform model, NO production was produced equally within the parenchymal domain. In the regional model, NO production within 50 µm was set to be 3.8 fold greater than the tissue further than 50 µm to mimic the increased perivascular density of nNOS neurons close to the vessel^32^. In the proximal model, all NO production in the parenchyma was restricted to within 2 µm of the arteriole wall. NO degradation in the parenchyma was dependent on the NO, oxygen, and cell concentration and expressed using the following equation^73^:

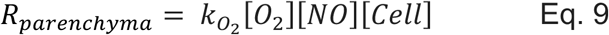

Where k_O2_ = 5.38 • 10^-10^ M^-1^s^-1^(cell/ml)^-1^ (**Table 1**) and the density of cellular sinks in the tissue ([Cell]) was chosen to be 10^Ñ^ cell/ml, as was previously used for NO diffusion modeling in parenchymal tissue^206^. Note that the degradation rate of NO in the parenchyma was not uniform throughout the tissue because the oxygen content of the parenchyma changes with distance from the arteriole. Because the rate of NO degradation is proportional to the oxygen content of the tissue (**Eq. 9**), the oxygen rich region of the parenchyma near the arteriole will have a higher degradation rate of NO than distant from the arteriole where the oxygen concentrations fall off. For all of the simulations presented with the exception of those in Figure 7, the oxygen concentrations changed slowly enough in time that they could be assumed to be at steady state. This allowed us to use the Krogh model, as it gave oxygen profiles identical to full simulations of diffusion of oxygen using Fick’s equations with little computational overhead. However, for simulations where rapid and large manipulations of oxygen tension were performed (**Figure 7**), we simulated the diffusion and consumption of oxygen in the parenchyma (**Eq. 3**). Within the NO producing region of the parenchyma, NO production/degradation was accounted for with a production rate P_NO_(t).

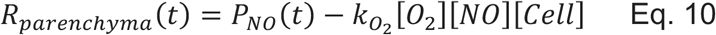

For steady-state simulations, P_NO_(t) was a constant production rate that was parametrically varied. For time-dependent simulations, P_NO_(t) was modified to be a pulse of increased NO production or white noise centered around the EC_50_ for guanylyl cyclase activity (8.9 nM).

### Simulating diffusion in a deforming domain

The deformation of the RBC core and the cell free layer during vasodilation and constriction were modeled with a linear displacement of the finite element mesh. The deformation of the parenchymal tissue, smooth muscle, and endothelial cell layer was modeled by linear elasticity^213^ with a Poisson ratio of 0.45. The spatial gradients in the diffusion equation in all the domains were transformed into gradients in deforming domains using the arbitrary Lagrangian-Eulerian (ALE) method^214^. The total parenchymal NO production rate was held constant during the slight deformation of the surrounding tissue due to changes in arteriole diameter. Because the smooth muscle was modeled as nearly incompressible and its cross-sectional area did not appreciably change, vasodilation reduced its thickness such that the distal boundary of the smooth muscle became closer to the lumen. Vasodilation also displaced the tissue radially outward and the displacement was taken into account when comparing points in the tissue at different dilation states.

### Power Spectrum Calculation

We investigated the preferred frequency of vasodynamics in the model by introducing a white Gaussian noise production rate of NO (30 Hz, low pass filtered < 2 Hz, 25 minute duration) in the parenchyma within 2 µm of the arteriole wall (proximal model). NO production was initially set such that GC activity in the smooth muscle was at EC_50_ (8.9 nM) and the variance from a white Gaussian noise change in NO production was chosen such that there was no change in vessel diameter exceeding ±5%. Vessel sensitivity was set to 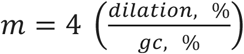 unless otherwise indicated. The power spectral density was calculated from the arteriole response in the model using the Chronux toolbox version 2.11 (http://chronux.org, function: mtspectrumc). We used 101 averages for a frequency resolution of 0.067 Hz.

### Calculation of the hemodynamic response function

The relationship between neural activity and vessel dynamics is often considered a linear time-invariant (LTI) system^211, 215, 216^ which allows for the hemodynamic response function to be calculated numerically using the relationship

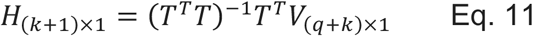

Where H is the HRF, V is the vascular response, and T is a Toeplitz matrix of size (q+k)× (k+1), containing measurements of normalized neural activity (n)^131^.

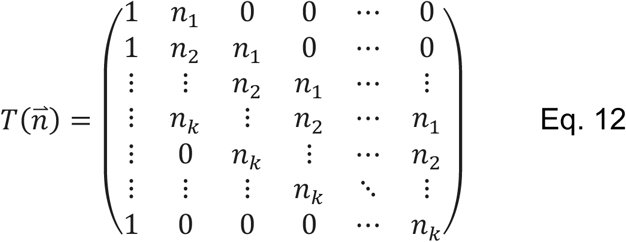

Note that this method makes no assumptions about the shape of the HRF. To evaluate the HRF produced in the model we performed the same calculation using a NO production rate (n) in place of neural activity, where n was white Gaussian noise.

### Estimating perivascular mitochondrial inhibition

Although NO dilates arteries, resulting in increased blood flow and O_2_ delivery to the tissue, it can also compete with O_2_ at the mitochondrial cytochrome c oxidase (CcO) to inhibit aerobic respiration and facilitate the generation of free radicals^183, 217^. Under physiologic conditions, inhibition of CcO by NO is minimal and reversible^56, 218–220^ but under conditions of high NO and/or low O_2_, CcO can be permanently inhibited^119^. Permanent inhibition of CcO occurs at nominal NO and O_2_ concentrations of 1000 nM and 130 μM, respectively^119^ which is equivalent to 12.5% CcO activity using a competitive model of inhibition:

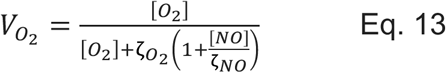

Here, V_O2_ is the fractional activity of CcO, ζ_O2_ = 210, ζ_NO_ = 0.225, and [O_2_] and [NO] are the respective oxygen and NO concentrations, expressed in nM^54, 217, 218^. Because permanent inhibition of CcO is likely pathological (V_O2_ ≤ 12.5%), it is unlikely that physiological NO-mediated NVC produces this combination of NO and O_2_ concentrations.

## Data Availability

Code used to generate the figures in this paper is available at: https://psu.box.com/v/Haselden-NO-Code

## Acknowledgements

This work was supported by R01EB021703 from the NIH to PJD. We thank Y. Kim and Y-T Wu for providing images of the brain vasculature in Figure 1.

---

**Figure 2–figure supplement 1.**
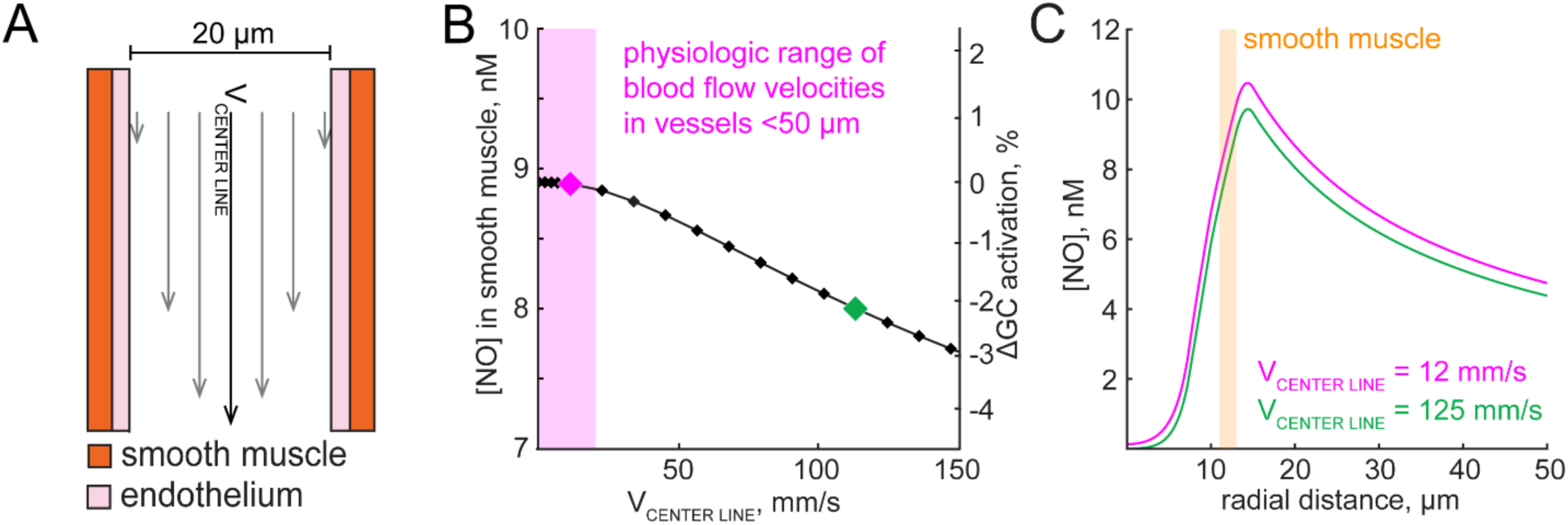
Convection has a negligible effect on perivascular NO concentrations. A) Relative velocity profile of blood flowing through an arteriole that is proportional to convective transport experienced by NO. B) Within physiologic blood flow velocities in a 20 μm diameter arteriole^222, 223^, convection causes a negligible change in [NO] (Δ0.1% GC activation). The pink diamond indicates a physiologic flow in a 20 μm diameter arteriole, while the green diamond is approximately tenfold higher, comparable to the centerline velocity in a 200 μm diameter arteriole^223^. C) Perivascular [NO] profiles accounting for convection at physiologic (pink) and extreme (green) blood flow velocities. [NO] profiles when blood flow velocities are less than 20 mm/s is conserved. Pink and green data diamonds in B are plotted as perivascular [NO] profiles in C.

**Figure 4–figure supplement 1.**
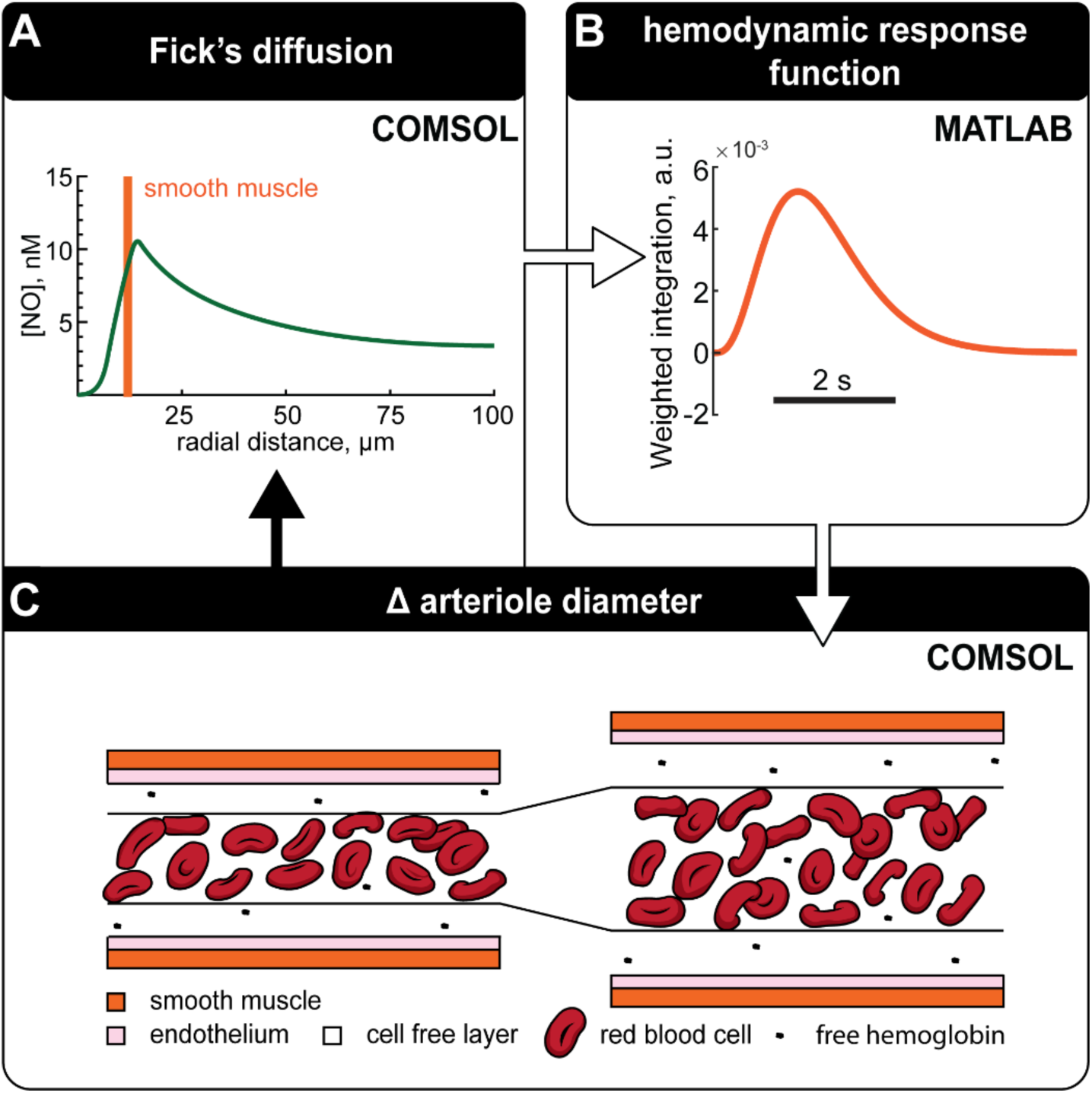
Coupled diffusion and deformation schematic. A) NO gradients surrounding an arteriole is evaluated using Fick’s equation. B) The recent history of NO levels in the smooth muscle domain is converted to GC activation and convolved with a kernel to account for the signaling cascade that converts GC activation into dilation. This kernel was chosen to match the temporal dynamics of neurally-evoked dilation. C) Arteriole diameter is adjusted depending on the output of the kernel with more or less NO corresponding to dilation and constriction respectively. Diffusion of NO can then be re-evaluated using the new arteriole geometry. Adjustments to arteriole diameter using this cycle are made at each time step.

**Figure 4–figure supplement 2.**
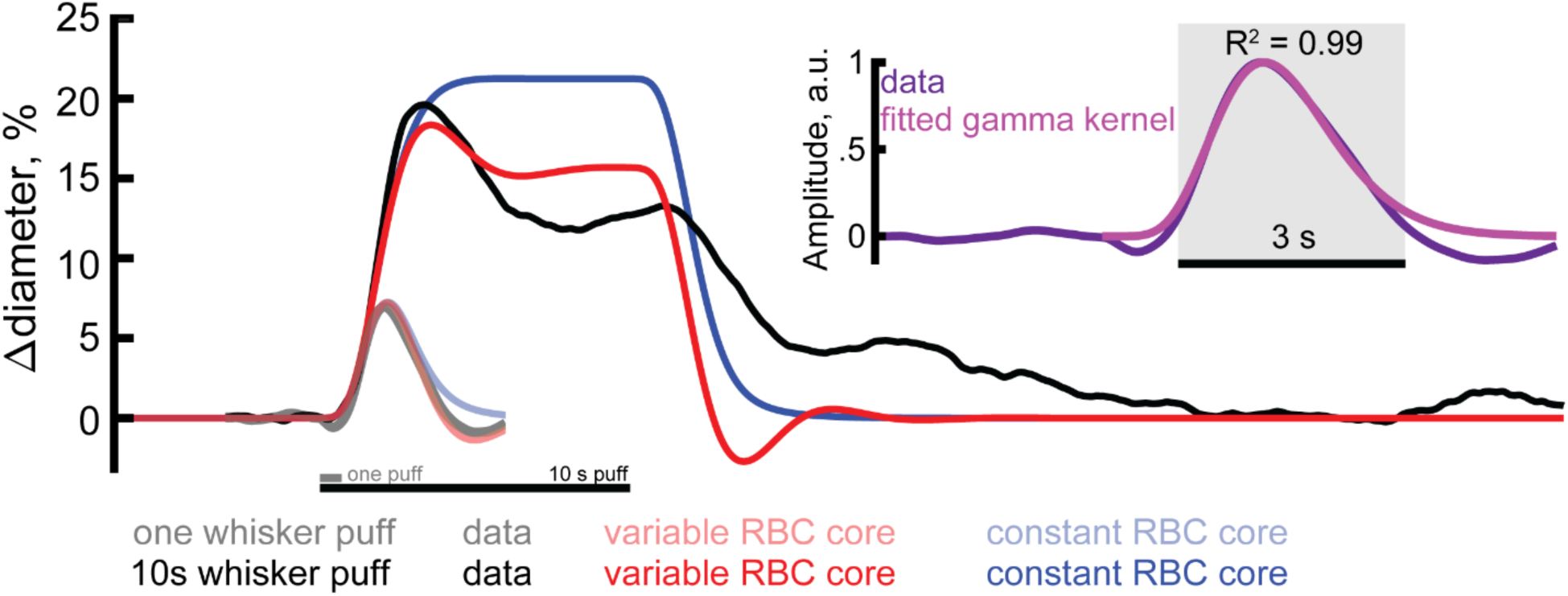
Model dynamics of allowing interplay of NO degradation and vasodilation. *In vivo* mouse surface arteriole diameters in the somatosensory cortex in response to a single and 10-second-long puff to the whiskers^1^ (black) were compared to the model with a variable RBC core (red) or constant RBC core (blue). The response kernel of the vessel was fitted to the positive component of the hemodynamic response function (HRF) from a single whisker puff (inset) and the slope of vessel sensitivity to NO, m, was set to 3. Allowing the degradation of NO to dynamically change with arteriole diameter imposed a post-stimulus undershoot that was not present when the RBC core diameter was held constant.

**Figure 5–figure supplement 1.**
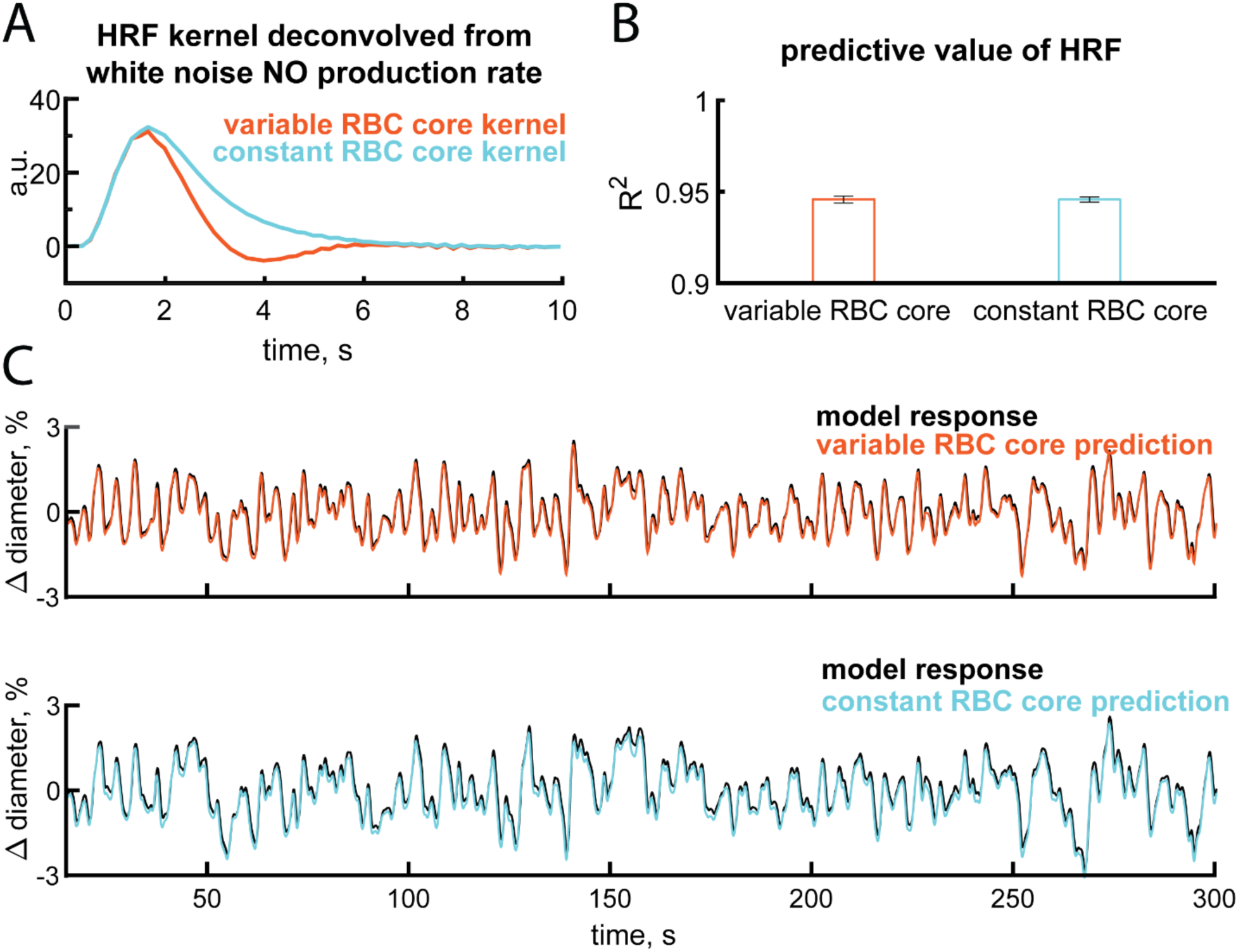
Coupling NO degradation to vessel size is a linear system. A) The variable and constant RBC core model HRF shown is deconvolved from 12 minutes of white noise NO production when 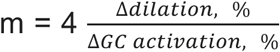. B) Convolution of the NO production rate with the kernel has a high predictive value for estimating the arteriole response for the remaining 12 minutes of the trial. R^2^ values shown are from m = 1,2,3,4, and 5. C) Example 300 seconds of data showing the difference between the response in the model and an approximation using the kernels shown in A. Example shown for m = 4.

**Figure 6–figure supplement 1.**
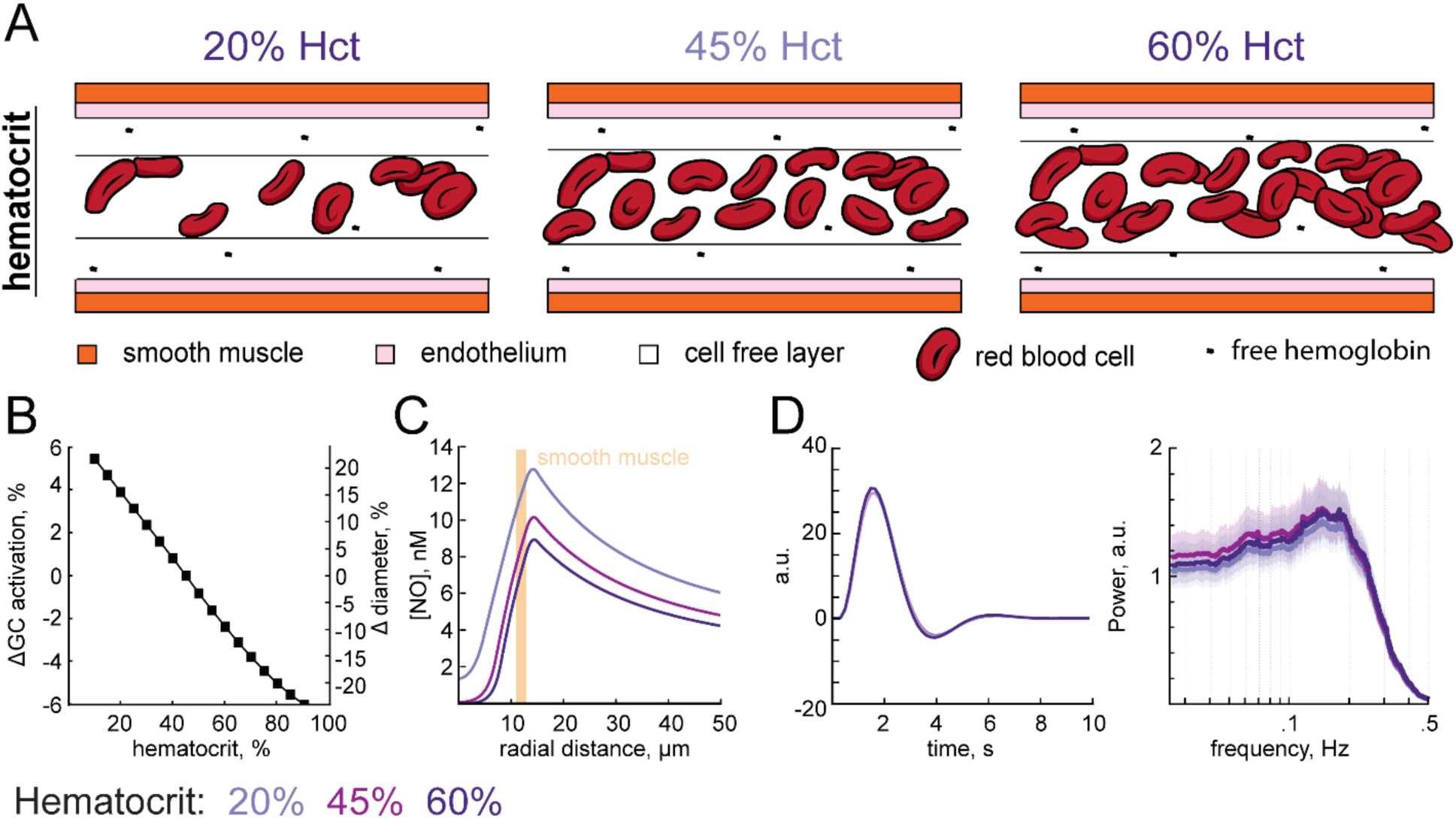
Impact of hematocrit on vasodynamics. A) Increasing hematocrit increases the degradation of NO in the RBC core and reduces the size of the cell free layer. B) Baseline GC activation and arteriole diameter are predicted to decrease with increasing hematocrit and increase with decreasing hematocrit. C) Perivascular NO increases with low hematocrit and decreases with elevated hematocrit. The location of the smooth muscle is indicated in orange. D) Changing hematocrit does not alter the hemodynamic response function or the frequency response of the vessels (E).

